# Competition co-immunoprecipitation reveals interactors of the chloroplast CPN60 chaperonin machinery

**DOI:** 10.1101/2023.01.05.522938

**Authors:** Fabian Ries, Heinrich Lukas Weil, Claudia Herkt, Timo Mühlhaus, Frederik Sommer, Michael Schroda, Felix Willmund

## Abstract

The functionality of essential metabolic processes in chloroplasts depends on a balanced integration of nuclear-and chloroplast-encoded polypeptides into the plastid’s proteome. The chloroplast chaperonin machinery is an essential player in chloroplast protein folding with a more intricate structure and subunit composition compared to the orthologous GroEL/ES chaperonin of *Escherichia coli*. However, its exact role in chloroplasts remains obscure, mainly because of a very limited knowledge about the folded substrates. We employed the competition immunoprecipitation method for the identification of the chaperonin’s substrates in *Chlamydomonas reinhardtii*. Co-immunoprecipitation of the target complex in the presence of increasing amounts of isotope-labelled competitor epitope and subsequent mass spectrometry analysis specifically allowed to distinguish true interactors from unspecifically co-precipitated proteins. Besides known substrates such as RbcL, we revealed numerous new substrates with high confidence. Identified substrate proteins differ from bulk chloroplast proteins by a higher content of beta-sheets, lower alpha-helical content and increased aggregation propensity. Immunoprecipitations performed with a subunit of the co-chaperonin lid revealed the ClpP protease as a specific partner complex, with altered interactions during heat stress, pointing to a close collaboration of these machineries to maintain protein homeostasis in the chloroplast.

## INTRODUCTION

The correct folding of proteins is a prerequisite for their biological activity and is predetermined by the amino acid sequence (Anfinsen, 1973). This intricate process is guided by a cascade of factors that promote folding, maturation, and complex formation within the crowded cellular environment for preventing misfolding and aggregation. This is achieved by a structurally versatile group of molecular chaperones that act at various stages during a protein’s life (Balchin et al., 2016). Of those, members of the Hsp60 chaperonin family form large barrel-shaped complexes that can encapsulate their client proteins in the so-called *Anfinsen cage*, an unperturbed environment where folding is facilitated (Saibil et al., 1993). Bacterial (GroEL), mitochondrial (Hsp60) and plastid (Cpn60) chaperonins belong to the group I chaperonin family and function together with the dome-shaped co-chaperonin GroES in bacteria, Hsp10 in mitochondria, and Cpn10/20 in plastids (Trösch et al., 2015). All group I chaperonins represent cylindrical complexes containing two stacked rings, composed of seven subunits per ring (Hayer-Hartl et al., 2016). In bacteria and mitochondria, chaperonins and their respective co-chaperonins are each formed by a single protein isoform. In contrast, different isoforms of Cpn60 and Cpn10/20 exist for the chloroplast chaperonin machinery and functional complexes accumulate in heteromeric compositions (Tsai et al., 2012, Trösch et al., 2015, Zhao et al., 2019, Vitlin Gruber et al., 2013). The unicellular green alga *Chlamydomonas reinhardtii* (Chlamydomonas herein) contains three Cpn60 isoforms (CPN60A of the alpha subgroup, CPN60B1 and CPN60B2 of the beta subgroup), which share only 50% sequence identity between the alpha and beta forms. Recent structural analysis of the Chlamydomonas CPN60 complex revealed a ratio ranging between 5:3:6 and 6:2:6 for CPN60A:B1:B2 (Zhao et al., 2019). Despite the heterogenic subunit composition of chloroplast chaperonins, the overall architecture and function appears to be conserved between bacteria and chloroplasts (Zhao et al., 2019). In addition, three isoforms (CPN11, CPN20 and CPN23) of the co-chaperonin are found (Schroda, 2004, Trösch et al., 2015). CPN20 and CPN23 comprise two GroES domains, which are fused in tandem and constitute the lid complex in a 1:2:1 stoichiometry for CPN11:CPN20:CPN23 (Tsai et al., 2012, Weiss et al., 2009, Zhao et al., 2019).

The chaperonin-mediated folding cycle is driven by ATP binding, hydrolysis, and release concerted with co-chaperonin association and dissociation. For GroEL, it was shown that cooperative ATP binding to the *cis* ring concomitant with substrate binding induces first structural changes resulting in tilting and elevation of the apical domains (Clare et al., 2012). The binding of a substrate onto these surfaces may also trigger its stretching, which causes unfolding of trapped substrate conformations. At this state, the co-chaperonin GroES closes the folding cage by binding to exposed hydrophobic apical sites of GroEL (Horwich and Fenton, 2020). GroES binding triggers a 100° clockwise twist of the apical GroEL domains, hence detaching the substrate off the GroEL walls and exposing them to a now hydrophilic chamber. The subsequent hydrolysis of ATP takes ∼10 seconds. Substrate and ATP binding to the opposite *trans* ring then causes GroES detachment from the complex and release of the folded substrate from the *cis* ring (Clare et al., 2012). In *E. coli*, the identified GroEL/ES substrates comprise approximately 10% of the entire proteome (Houry et al., 1999, Ewalt et al., 1997, Kerner et al., 2005). These substrates are enriched for (β⍺)_8_ TIM (Triose-phosphate isomerase) barrel folds and occupy one third of the GroEL/ES capacity (Kerner et al., 2005). The identified ∼250 substrate proteins were subdivided into four classes, based on their increasing dependence on GroEL/ES for folding (Niwa et al., 2016, Kerner et al., 2005). The strict GroEL dependence of ∼50 substrate proteins is probably the reason why GroEL/ES is indispensable for viability of *E. coli* under all conditions (Horwich et al., 1993, Fayet et al., 1989).

In chloroplasts, it was early discovered that maturation of RbcL, the large subunit of the Ribulose-1,5-Bisphosphate Carboxylase/Oxygenase (Rubisco), depends on the chaperonin (Barraclough and Ellis, 1980, Horwich and Fenton, 2020, Mizohata et al., 2002). Surprisingly, only few other plastidic folding substrates are known to date, but there is accumulating evidence that the chaperone is involved in the folding of imported and chloroplast-encoded nascent polypeptides (Trösch et al., 2015, Ries et al., 2020). We have recently reported that the chaperonin machinery associates with translating plastidic ribosomes and that this binding increases during heat acclimation, suggesting that the chaperonin binding of nascent proteins starts co-translationally minimizing the risk of misfolding (Westrich et al., 2021, Trösch et al., 2022). Interestingly, CPN60 seems also involved in guiding imported thylakoid membrane proteins through the stroma, as shown for Plsp1, the Plastidic type I signal peptidase 1 (Klasek et al., 2020).

For a better understanding of the functions of the chaperonin machinery in chloroplasts, we aimed to extend the knowledge about the substrate spectrum. We employed competition co-immunoprecipitation (competition co-IP) coupled with mass spectrometry as an approach that is well suited to distinguish true interactors from unspecifically co-precipitating contaminants during IP (Sommer et al., 2014). By determining the direct correlation how the target protein and its binding partner is titrated away via the competing epitope, we identified 38 high-confidence substrates of the chloroplast chaperonin. Some of these novel chaperonin substrates are highly abundant client proteins, highlighting the essential function of the chloroplast chaperonin. We further gathered support for the recent discovery of a close inter-connection between the chaperonin lid and the ClpP-protease, and we show increased interaction during heat exposure, suggesting enhanced interaction during situations of challenged protein homeostasis.

## RESULTS

### Establishing competition co-immunoprecipitation for CPN20 and CPN60A

The chloroplast chaperonin system, like its bacterial counterpart, is thought to encapsulate unfolded proteins to promote their folding (Figure 1a) (Trösch et al., 2015, Horwich and Fenton, 2020, Hayer-Hartl et al., 2016). Identifying substrates of this chaperonin system via co-immunoprecipitation and subsequent mass-spectrometry (co-IP-MS) depends on a reliable method to distinguish true interactors from unspecific contaminants, with the latter being co-purified due to unspecific binding to the applied beads and antibodies (ten Have et al., 2011, Markham et al., 2007). Competition co-IP was previously reported to be well suited for such a distinction (Sommer et al., 2014). Here, an inactivated derivate of the antigen, which cannot bind to its native protein partners, is added in a serial dilution to cell lysates before co-IP. The antigen competes with the target protein for antibody binding and, therefore, increased concentrations of this competitor result in the depletion of the purified target protein and its binding partners, while unspecific binders remain unaffected (Figure 1b).

**Figure 1:**
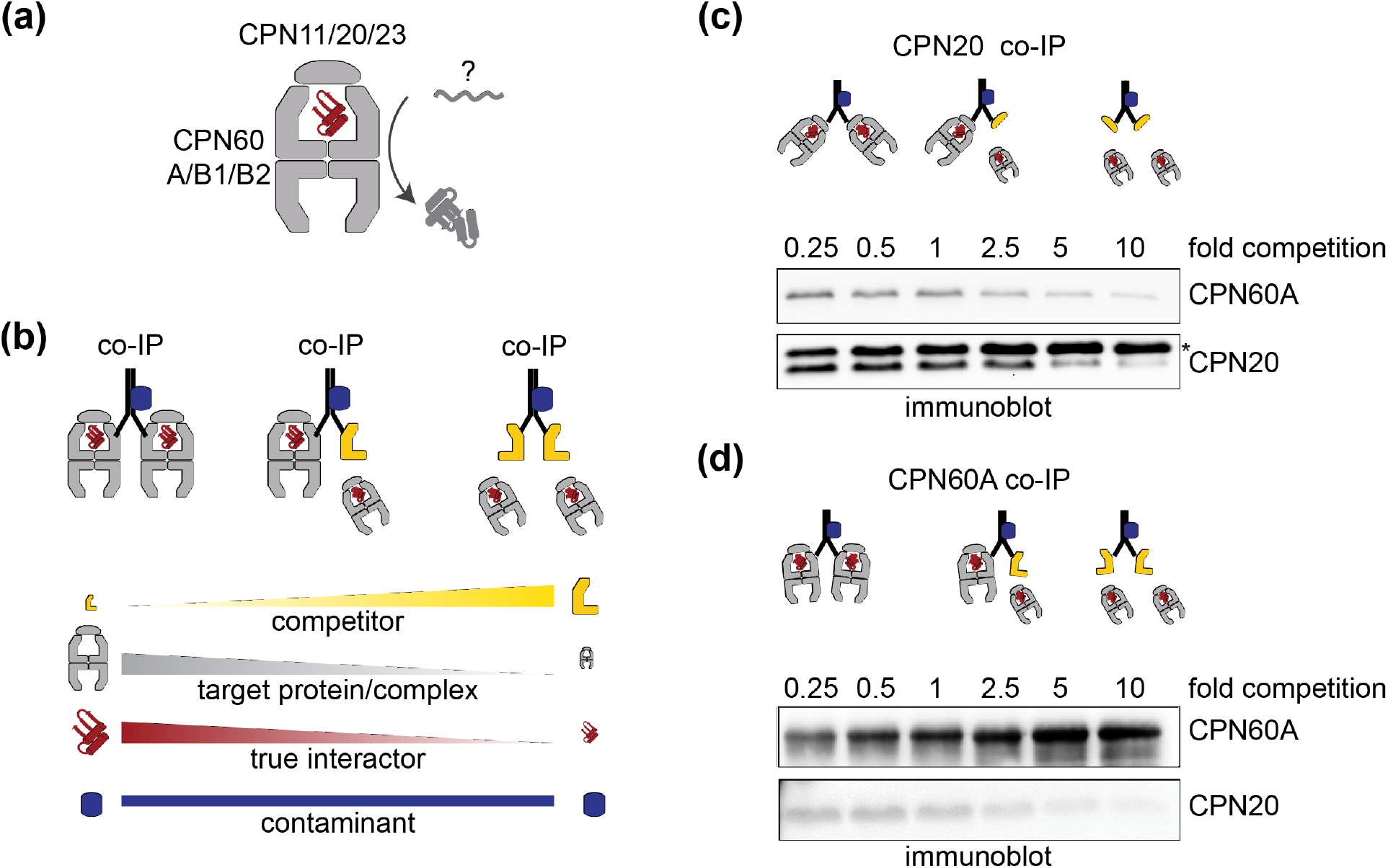
Setup of competition co-immunoprecipitation for CPN20 and CPN60A. (a) Schematic view representing protein folding via the chaperonin machinery consisting of the CPN60A/B1/B2 subunits and the co-chaperonin lid components CPN11/20/23, for which the substrates are largely unknown in the chloroplast (labeled “?”). (b) Setup of competition co-immunoprecipitation (co-IP) and the expected protein abundance changes in eluates with increasing competitor concentrations (yellow symbol). (c) Increased CPN20 competitor amounts reduce the abundance of complex partner CPN60A in precipitates. The amount of competitor added relative to endogenous CPN20 in the lysate (fold competition) is indicated. The CPN20 competitor harbors a hexa-his-tag (indicated with an asterisk) and therefore has a larger apparent molecular mass than endogenous CPN20. (d) Increased CPN60A competitor amounts reduce the abundance of complex partner CPN20 in precipitates. Untagged competitor CPN60A migrates with the same apparent molecular mass as endogenous CPN60A and therefore masks decreasing CPN60A levels.

We aimed to identify interactors or substrate proteins of the chaperonin machinery by targeting a subunit of the CPN60 barrel complex as well as by targeting a subunit of the lid complex. The latter allows identifying enclosed substrates by purifying lid-bound chaperonin complexes (Kerner et al., 2005). For the barrel, CPN60A appeared best suited since it is abundantly present in CPN60 complexes of Chlamydomonas cells and most divergent from the other two subunits (CPN60B1 and CPN60B2) (Zhao et al., 2019). In addition, we used an antibody, raised against a section covering only the C-terminal 124 amino acids, which still provides sufficient epitope at the outside of the CPN60 barrel (Figure S1) to lower cross-reactivity with the other subunits. Full-length and mature CPN60A, missing the 33 amino acids of the predicted chloroplast transit peptide (Figure S1a), was used as competitor. For enriching the lid-bound complex, an antibody was raised against CPN20, which is the predominant subunit in the co-chaperonin complex (Zhao et al., 2019, Tsai et al., 2012). The mature CPN20 protein, lacking the 22 amino acids of the predicted chloroplast transit peptide, was used as competitor. Both competitors were heterologously expressed in *E. coli* grown on ^15^NH_4_Cl, in order to distinguish the isotope-labelled competitor from the respective unlabelled target protein in immunoprecipitates from Chlamydomonas cell extracts by mass spectrometry (Sommer et al., 2014). Competitor proteins were not inactivated since we considered an exchange with preformed protein complexes in cell lysates as unlikely in ATP-depleted samples (Bai et al., 2015).

We first estimated the protein abundance of CPN20 and CPN60A in Chlamydomonas cells. Based on quantitative immunoblotting of cell lysate samples and defined amounts of full-length, purified CPN20 and CPN60 proteins, we estimated that CPN20 and CPN60A represent about 0.17 ± 0.06% and 0.1 ± 0.04% of total cellular proteins, respectively (Figure S2), consistent with previous abundance ranking based on shotgun proteomics data that showed a higher abundance of CPN20 compared with CPN60A (Schroda et al., 2015). To ensure competition between the target protein and the purified epitope for antibody binding sites, it is required to employ sub-stoichiometric amounts of antibodies in the IP relative to the target protein. In a lysate derived from 5×10^8^ Chlamydomonas cells, a range of 5 to 50 μg affinity-purified and immobilized antibody yielded a linear increase of precipitated CPN20 protein and the known substrate RbcL, while CPN20 was only mildly depleted from the sample (Figure S3). Thus, 25 μg CPN20 and CPN60 antibodies per lysate from 5×10^8^ Chlamydomonas cells were used for all future experiments. To stabilize protein interactions, samples were chemically crosslinked with 2 mM dithio-bis(succinimidyl propionate) (DSP) upon lysis. A range of 0.25- to 10-fold excess (0.25, 0.5, 1, 2.5, 5, 10-fold) of competitor, relative to endogenous CPN20 or CPN60A protein, resulted in a linear decrease of co-precipitation of the respective complex partner of CPN60A or CPN20, indicating that this selected range was well suited for the competition series (Figure 1c and 1d).

### Identification of CPN20 and CPN60 interactors by mass spectrometry

For the identification of chaperonin interactors or substrates via mass spectrometry, three independent co-IPs were performed with antibodies targeted against CPN20 or CPN60A, respectively. Increasing amounts of ^15^N-labelled competitor protein were added to the IPs as described above. Importantly, peptide ion intensity-based quantification of the CPN20 and CPN60A ^15^N competitor versus ^14^N endogenous target protein resulted in ratios matching the calculated ones, particularly for CPN60A (Figure 2a). Furthermore, successful competition was validated by a similar decline of ^14^N target proteins (CPN20 and CPN60A) and their known complex partners in the precipitates (CPN11/23 and CPN60B1/B2, respectively) (Figures 2b and 2c).

**Figure 2:**
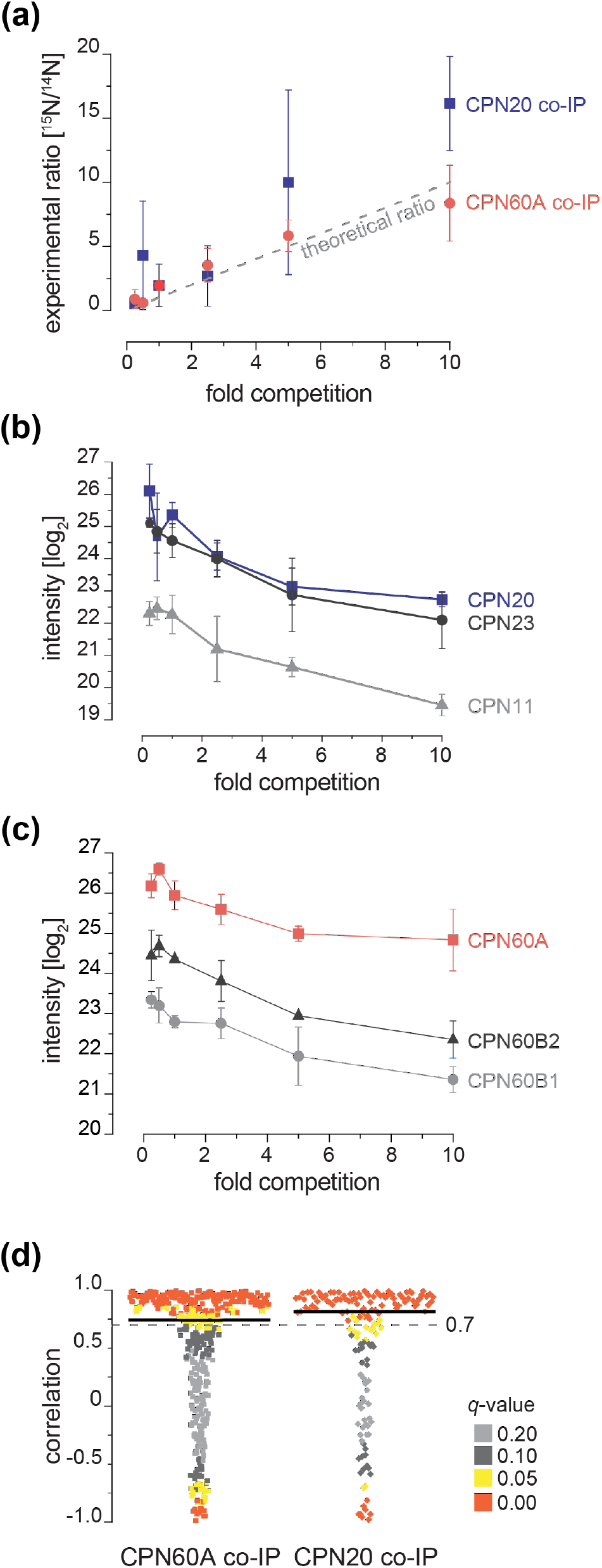
Mass spectrometry-based quantification of competition effects. (a) Measured ratios of ^15^N competitor to ^14^N target protein plotted against the ratio of competitor to endogenous target (“fold competition”). The theoretical ratio is given as grey dashed line. (b) and (c) Chaperonin subunits co-precipitated with CPN20 and CPN60A, respectively. Data points indicate mean values of three independent experiments with independently grown cells, error bars denote standard deviations. (d) Distribution of correlation coefficients for all co-precipitating proteins in CPN20 and CPN60 competition co-IPs, relative to their respective target protein. The color code indicates *q*-values of the respective proteins. Candidate proteins that were considered for further analyses had correlation coefficients >0.7 and a *q*-value <0.05.

For each dataset, only proteins were considered that were identified in precipitates obtained with at least five of the six competitor concentrations (Supporting Dataset). *PEARSON* correlations were calculated to determine the proteins that displayed a similar pattern of decreasing peptide ion intensities with increasing competitor concentrations as determined for the target protein (see Experimental Procedures). Of the 164 identified proteins in the CPN20 co-IPs and the 459 proteins in the CPN60A co-IPs, 90 and 190, respectively, showed *PEARSON* correlation values greater 0.7 and *q*-values smaller than 0.05 (red data points, Figure 2D) (Supporting Dataset). The different number of putative interactors in the CPN20 and CPN60A could be the result of different depths in the mass-spectrometry measurements, but it could also result from a higher likelihood of enriching interactors from the chaperonin’s folding chamber. Nevertheless, the known substrate RbcL correlated well with *r*-values of 0.95 (CPN60A) and 0.99 (CPN20), comparable with hitherto unknown chaperonin substrates such as GAPA1, SHK1 or AtpA (Figure 3a and 3b). In contrast, the uncharacterized protein Cre16.g688302 and HSP70A can be designated as contaminants based on low *r*-values and high *q*-values (Figure 3c). Remarkably, we identified seven out of eight subunits of the plastid-localized ClpP protease complex, which was specifically identified in the CPN20 co-IPs, except for chloroplast-encoded ClpP1 being also enriched in the CPN60 co-IP (Table 1). This is consistent with previous reports that the co-chaperonin complex co-purifies with the ClpP protease when isolated from Chlamydomonas lysates (Wang et al., 2021), indicating that proteins with high *r*- and low *q*-values are good candidates for chaperonin substrate or partner proteins.

**Table 1:**
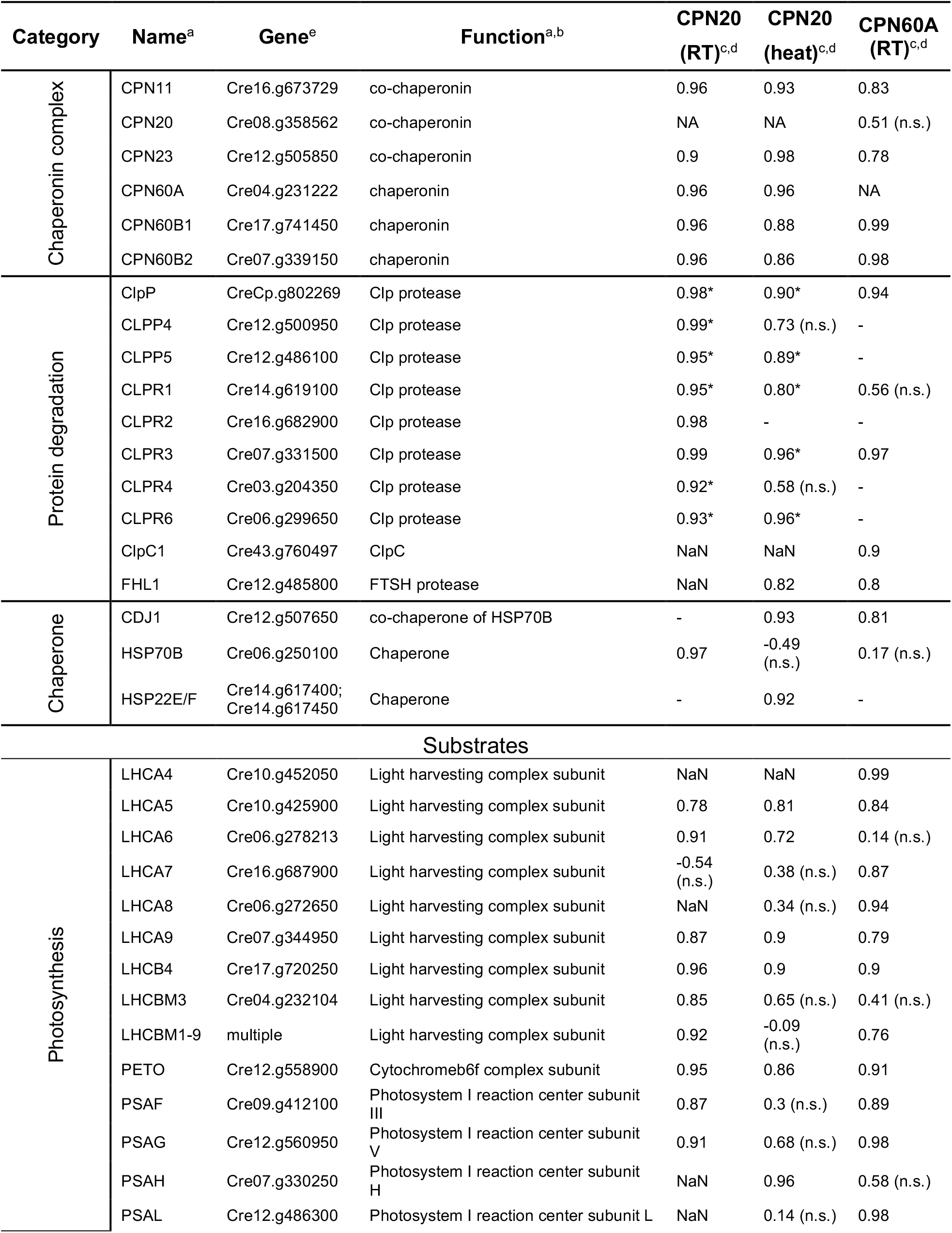
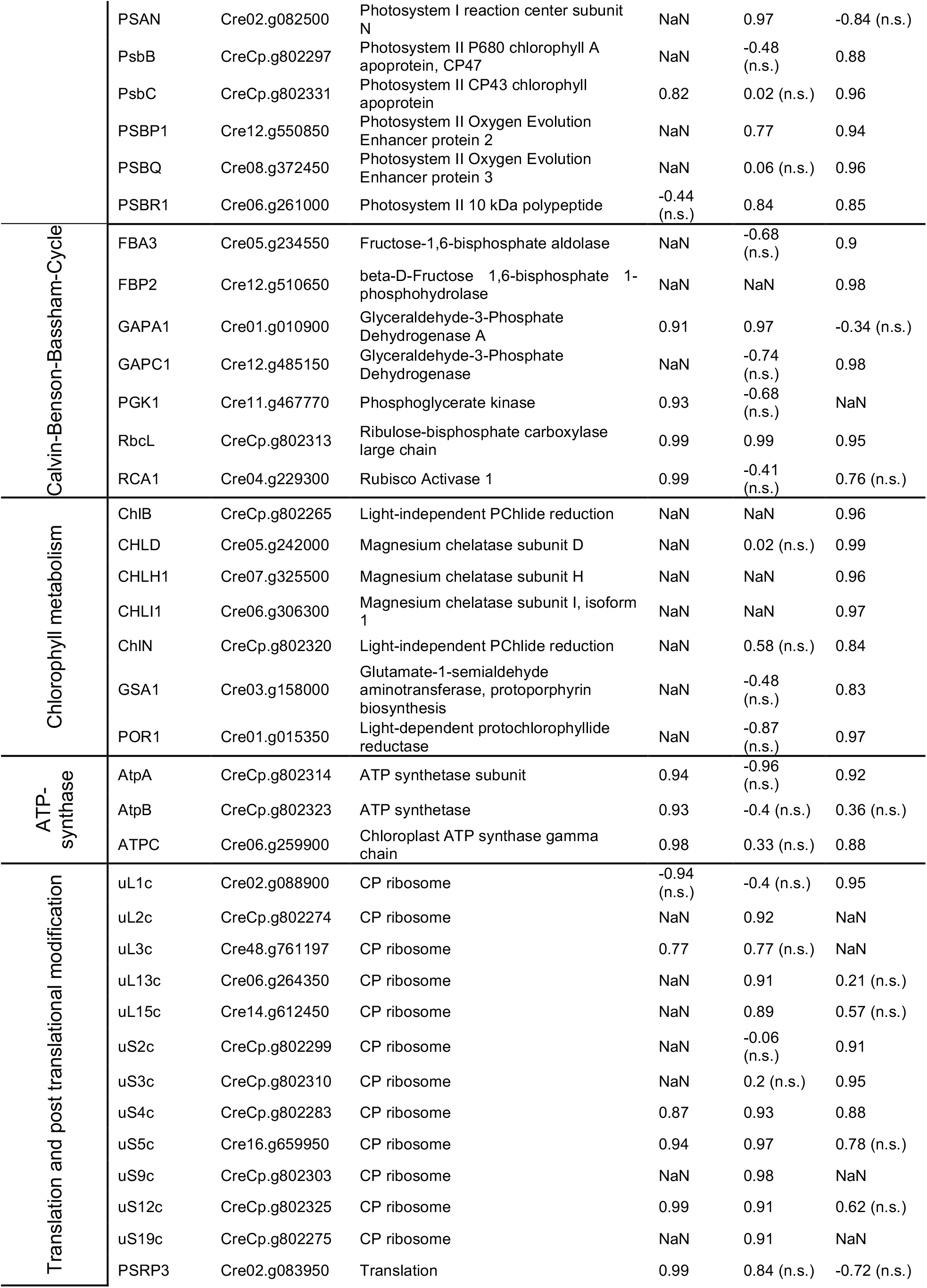
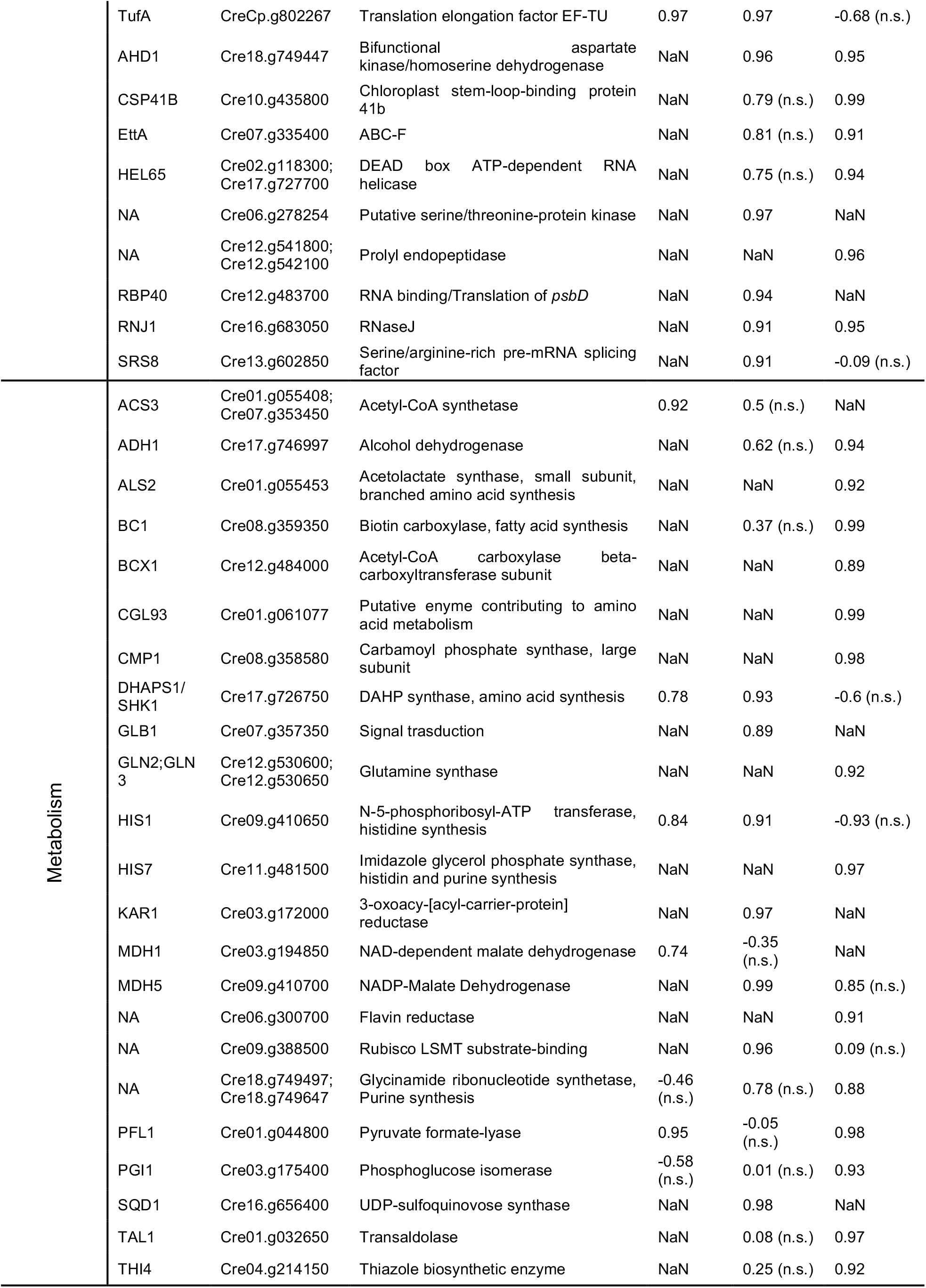
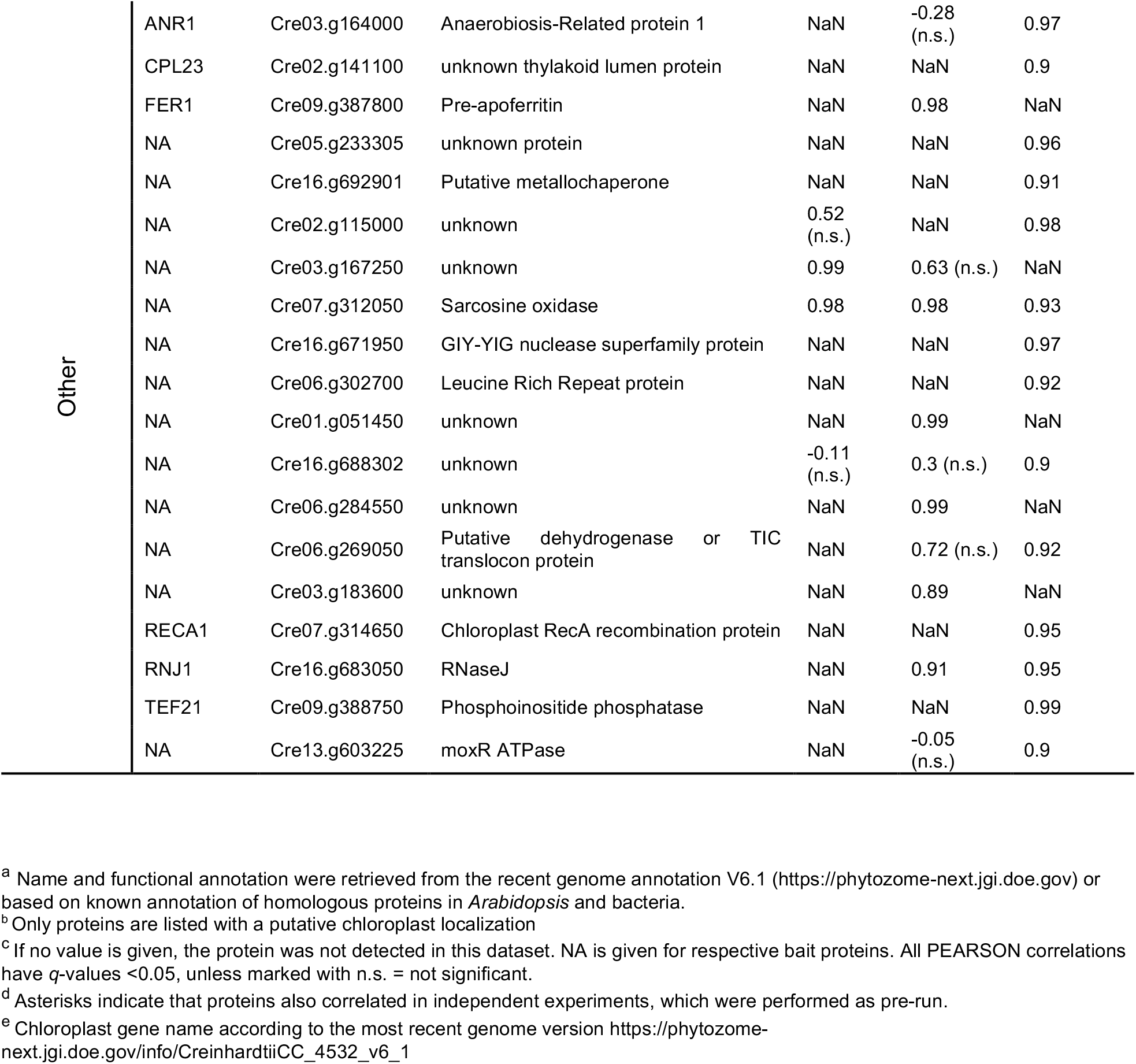
Interactors and substrates of CPN20 and CPN60A. For the complete list, see Supporting Dataset

**Figure 3:**
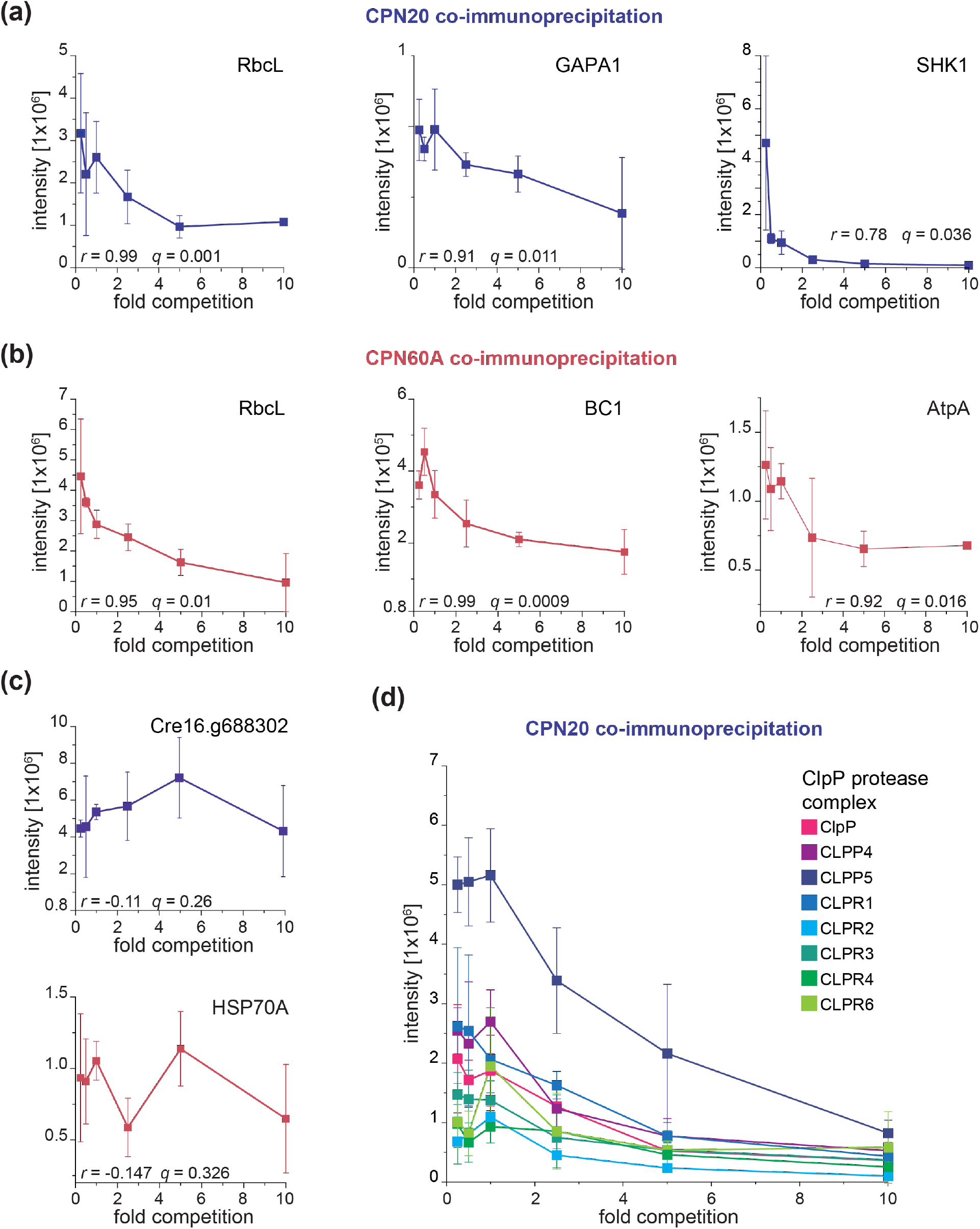
Correlation of abundances of proteins identified in competition-IPs with their CPN20 and CPN60 targets. Summed peptide ion intensities of proteins measured in at least five of six competition-IPs with CPN20 and CPN60A are plotted versus the ratio of added competitor to endogenous protein. (a) High-confidence substrates of CPN20. (b) High-confidence substrates of CPN60A. (c) Exemplary plots for proteins that were assigned as contaminants based on low *r*- and high *q*-values. (d) Summed peptide ion intensities of catalytic (CLPP) and regulatory (CLPR) subunits of the plastidic ClpP protease in CPN20 co-IPs are plotted versus the ratio of added CPN20 competitor to endogenous CPN20. RbcL = Large subunit of Rubisco; GAPA1 = glyceraldehyde-3-phosphate dehydrogenase A subunit; SHK1 = 3-deoxy-D-arabino-heptulosonate 7-phosphate synthase 1; BC = Biotin carboxylase; AtpA = CF1 ATPase beta-subunit, HSP70A = cytosolic Heat-Shock-Protein A.

### Properties of the chaperonin substrates

Of the 90 and 190 proteins with significant *r*- and *q*-values in the CPN20 and CPN60A datasets, only 51% and 42%, respectively, have a predicted or known chloroplast localization, whereas most remaining proteins are cytosolic, and several binding proteins are mitochondrial (Figure 4a, Supporting Dataset). We assume that these proteins bind to the CPN60 machinery after cell lysis.

**Figure 4:**
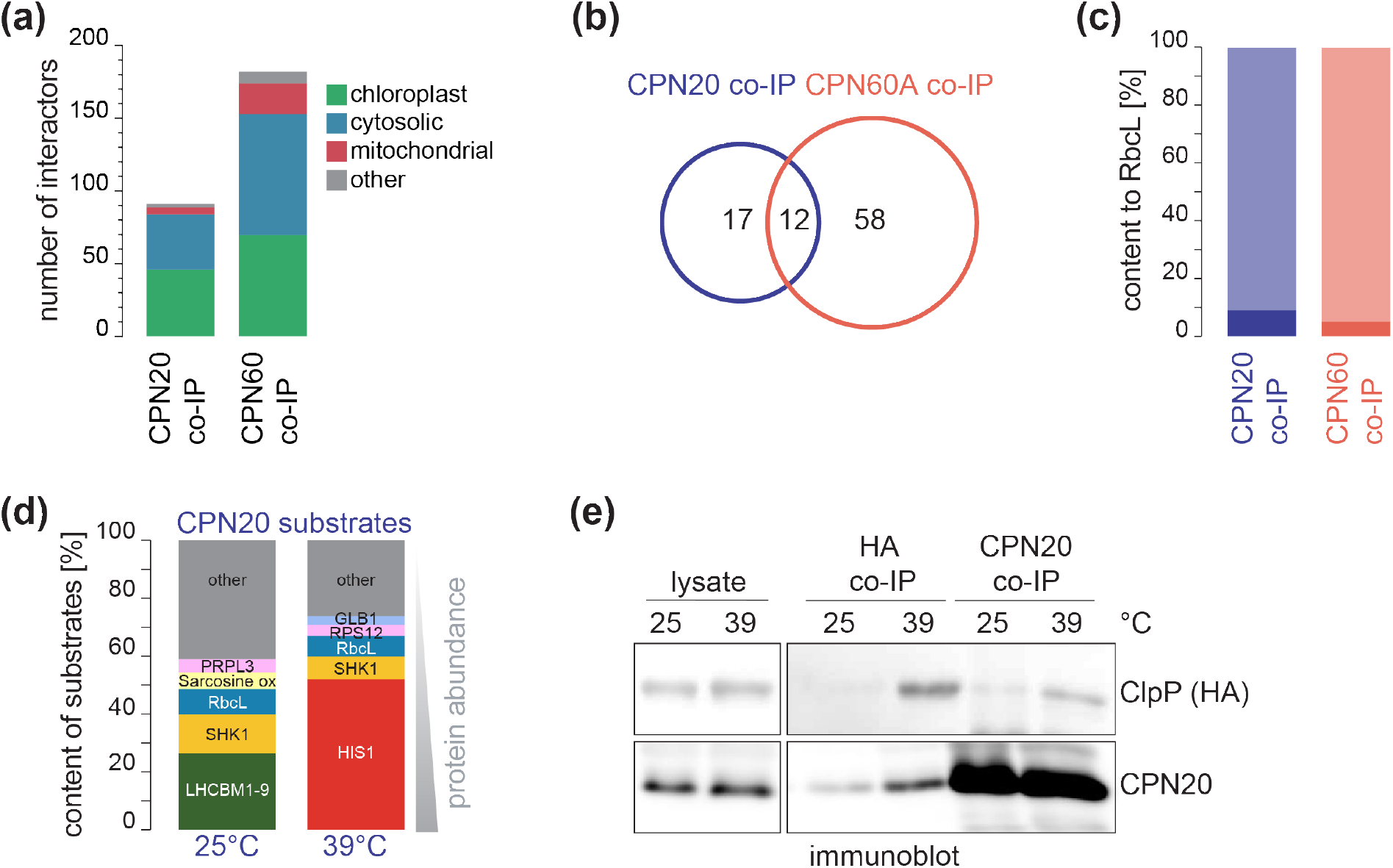
Properties of CPN20 and CPN60 substrates. (a) Subcellular localization of the proteins in the CPN20 and CPN60A datasets with correlations >0.7 and *q*-values <0.05. (b) Venn diagram representing the overlaps of chloroplast-localized substrate proteins of the CPN20 and CPN60A datasets (excluding ClpP complex subunits or the co-chaperonins themselves). (c) Bar graph comparing the fraction of RbcL (deep colored) in the co-precipitated chloroplast-localized substrates in CPN20 and CPN60A co-IPs. (d) Bar graph comparing substrates with a fraction larger than 5% (<5% summed in “other”) of chloroplast localized substrates in CPN20 co-IPs performed at 25°C and after a shift to 39°C for 30 min. (e) Immunoblot of co-IPs with antibodies against CPN20 and the HA epitope under the conditions described in (d) using a strain expressing ClpP with an HA-tag.

To further characterize the chaperonin substrates, only proteins with predicted chloroplast localization were considered and chaperonin or lid complex partners as well as the components of the ClpP complex were removed from the dataset. Of the remaining 29 and 70 high-confidence substrates in the CPN20 and CPN60A datasets, respectively, only twelve were found in both datasets (Figure 4b), suggesting that this substrate list is far from exhaustive. An explanation would be, that the prominent and highly abundant chaperonin substrate, RbcL, might tie up the vast folding capacity of the chaperonin machinery with limited capacity for other substrates. However, summed RbcL peptide intensities accounted only for 4-9% of total peptide intensities in the substrate data set (Figure 4c). Thus, it is unlikely that the identification of substrates was hindered by RbcL occupying most of the chaperonin’s folding capacity.

We further determined if the substrate spectrum is distinct under conditions of challenged protein homeostasis. To this end, the CPN20 co-IPs were also conducted from lysates of heat-exposed cells. Chlamydomonas cultures were shifted to 39°C for 30 min, a temperature which is still in the physiological range, causing only mild protein aggregation (Ries et al., 2017, Trösch et al., 2022, Rütgers et al., 2017). Interestingly, the prominent folding substrates of CPN20, such as the LHCs showed strongly reduced abundance within the substrates, while the faction of SHK1 and RbcL seemed reduced to half the intensity. The reduced substrate abundance goes in hand with our previous observation that *rbcL* translation is reduced under heat exposure (Trösch et al., 2022). Remarkably, the low abundant substrate HIS1 increased its intensity to 51% in the heat condition (Figure 4d, Table 1). In addition, we observed an increased interaction between the chaperonin lid complex and the ClpP machinery during heat exposure, indicating that this interaction might be stronger under conditions of challenged protein homeostasis (Figure 4e).

In agreement with other chaperonin systems, most of the substrates had a molecular weight below 60 kDa, which is considered as upper size limit for folding within the chaperonin’s cavity (Zhao et al., 2019) (Figure 5a and Figure S4). We further assayed the substrate proteins for an enriched occurrence of specific secondary structures in the datasets when compared to the total structural content of all chloroplast proteins. We found that proteins with high alpha-helical content are under-represented (Figure 5b), whereas proteins with high beta-sheet content are enriched (Figure 5c). In contrast, we found no significant enrichment regarding the physicochemical properties like amphiphilicity, hydrophobicity, coil-conformation, and isoelectric point (Figure S5). Using the TANGO algorithm, we estimated the aggregation potential of chloroplast proteins, based on the occurrence of aggregation-prone amino acid stretches within the amino acid sequence. Interestingly, the substrate proteins of the CPN20 and CPN60A dataset showed a significantly higher propensity for aggregation (Figure 5e). These properties showed a similar trend for the proteins found under heat conditions (Figure 5). Thus, a principal property of folding via the chaperoning complex could be the relative content of beta-sheet conformations and the potential for aggregation.

**Figure 5:**
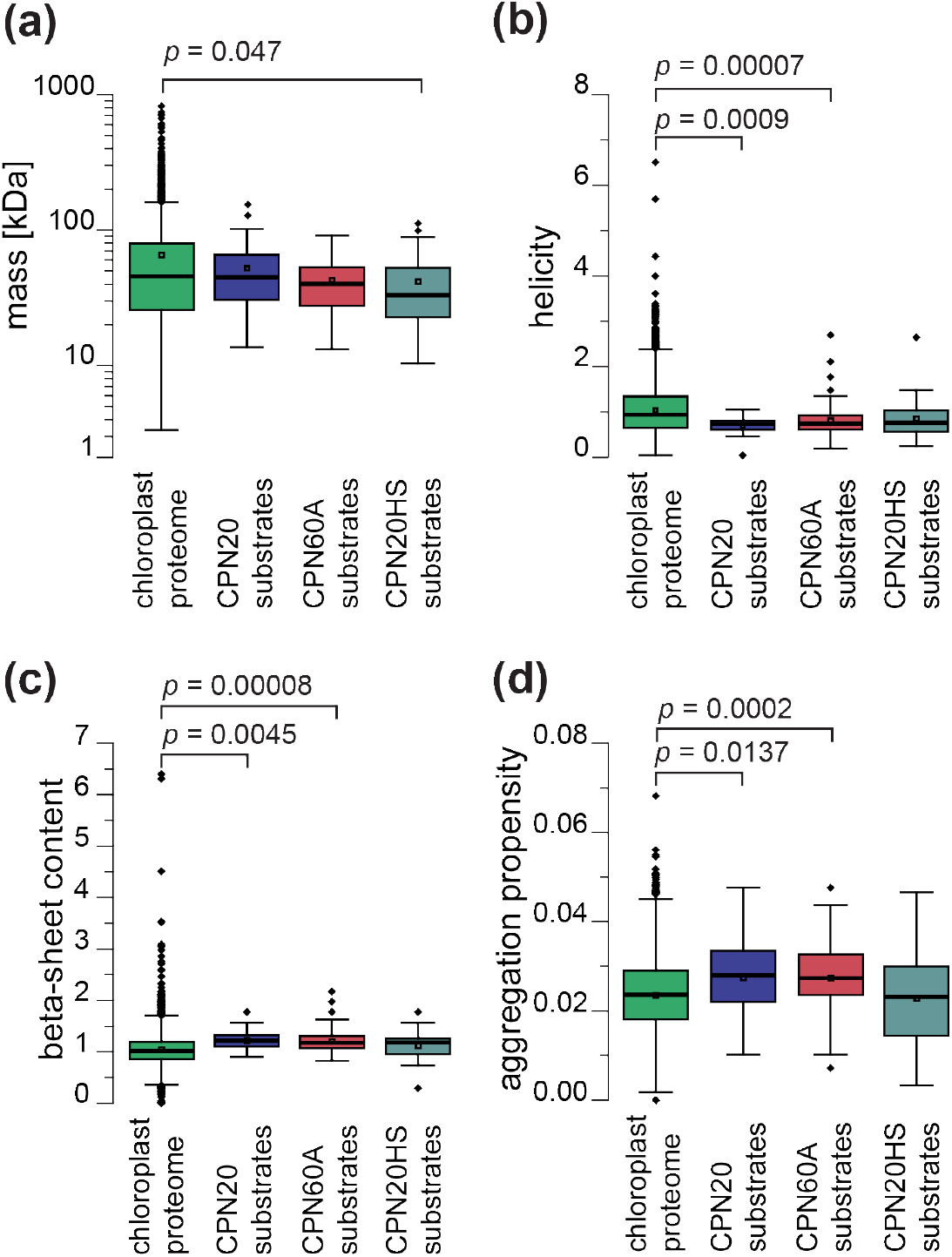
Properties of chaperonin substrates. Comparison of all chloroplast-localized proteins encoded by the Chlamydomonas genome (green), substrates of CPN20 (blue), CPN60A (red) identified under ambient conditions, and of CPN20 after a shift to 39°C for 30 min (turquoise). (a) Box plot comparing the distribution of molecular weights (based on complete protein sequences). (b) Boxplot representing the predicted alpha-helical and (c) beta-sheet content for proteins in the datasets. (d) Box plot showing values of predicted aggregation propensities as determined by TANGO, resulting in the number of aggregation prone islands relative to polypeptide sequence length. Students *t-*test based *p*-values are given for the significant differences to the chloroplast proteome.

## DISCUSSION

### Chloroplast chaperonin substrates and their physicochemical properties

The chloroplast chaperonin system and its lid co-chaperonin play an important role in this organelle, as demonstrated by growth defects when individual CPN60 components were reduced (Suzuki et al., 2009, Feiz et al., 2012, Bai et al., 2015, Wu et al., 2020). This is consistent with the essential role of chaperonins in the eukaryotic cytosol and in bacteria, where they are involved in folding of at least 10% of the respective proteome (Yam et al., 2008, Kerner et al., 2005, Chapman et al., 2006, Horwich and Fenton, 2020). However, it is not clear to date if the CPN60 machinery contributes to a similar level to protein folding in the chloroplast.

The most prominent folding substrate of CPN60 is the large subunit of Rubisco, RbcL, which also shows correlation values of ∼0.99/0.95 in our competition co-IP experiments with CPN20 and CPN60A (Table 1). Interestingly, comparison of substrate abundances in our datasets indicate, that RbcL is not the dominating substrate (with less than 10% abundance, Figure 4c), despite the high translation rates in chloroplasts of Chlamydomonas and an abundance of approximately 6% of the soluble proteome (Hammel et al., 2020, Hammel et al., 2018, Trösch et al., 2018). In the CPN60A dataset, GAPA1 is the most abundant high-confidence interactor (Supporting Dataset). GAPA1 (glyceraldehyde-3-phosphate dehydrogenase), the enzyme converting 1,3-biphosphoglyceric acid to glyceraldehyde 3-phosphate in the Calvin-Benson-Cycle in Chlamydomonas is forming a homo-tetramer (Avilan et al., 1997), in contrast to Arabidopsis where two isoforms form this complex (A_2_B_2_). Importantly, the bacterial ortholog G3P1 was also reported to partially depend on folding by GroEL/ES (Houry et al., 1999, Kerner et al., 2005). Previous *in vitro* experiments postulated a chaperonin dependence of the CF_1_ head subunits of the chloroplast ATP-synthase AtpA (⍺ subunit), AtpB (β subunit) and ATPC (*γ* subunit) for folding (Chen and Jagendorf, 1994, Mao et al., 2015). We found the AtpA and ATPC subunits in both the CPN20 and the CPN60 datasets, and AtpB only significant in the CPN20 dataset (Table 1), suggesting that these proteins require the chaperonin for *de novo* folding and assembly of the active CF_1_ core in chloroplasts (Mao et al., 2015). Interestingly, all three subunits are also part of the regulatory CES (Control by Epistasy of Synthesis) cascade in Chlamydomonas, which is essential for orchestrating the assembly of nucleus-derived (ATPC) and plastid-encoded subunits (AtpA and AtpB) (Drapier et al., 2007).

Despite the differences in subunit composition, the architecture of the chloroplast CPN60 complex and GroEL are surprisingly similar regarding the folding chamber (Zhao et al., 2019). The *Anfinsen cage* of GroEL, enclosed by GroES, is thought to comprise 175,000 Å^3^ (Xu et al., 1997) and thus limits the size of encapsulated proteins to a maximum molecular mass of approximately 60 kDa (Houry et al., 1999, Kerner et al., 2005, Hayer-Hartl et al., 2016). Consistently, most of our identified substrates were within a molecular mass range of 20-50 kDa (Figures 5a and Figure S4), allowing encapsulation for folding. Larger GroEL substrate proteins are found but these were postulated to be rather protected from aggregation instead of being actively folded (Chaudhuri et al., 2009). Interestingly, the enrichment of beta-sheet secondary structures and lower levels of alpha-helical conformations agrees with the properties of substrates of the cytosolic CCT/TRiC chaperonin (Yam et al., 2008), but these properties were not evidently enriched amongst substrates of GroEL/ES (Houry et al., 1999, Kerner et al., 2005). Proteins with beta strand conformations are thought to be more difficult to fold and to exhibit a higher aggregation propensity (Yam et al., 2008, Richardson and Richardson, 2002, Willmund et al., 2013), a property also shared amongst the chloroplast chaperonin substrates (Figure 5d). This goes in hand with previous reports, indicating that the structural properties of CPN60-dependent substrates are more complex and that chloroplast proteins cannot always be folded by the bacterial GroEL/ES system (Aigner et al., 2017, Chen and Jagendorf, 1994). Typical GroEL-dependent substrates are enriched in the TIM-barrel fold (Kerner et al., 2005). Due to limited structural information of chloroplast proteins, we were not able to globally investigate if these TIM barrel folds are also a characteristic feature of Chlamydomonas chaperonin substrates. However, RbcL and the aldolases FBA3 and TAL1 (Cre01.g032650 and Cre05.g234550), are examples for proteins with TIM-barrel fold (Table 1), indicating that such structural conformations may also require CPN60 for folding.

### Folding of nascent polypeptides

Most chloroplast-localized proteins are imported once synthesized in the cytosol. Several stromal chaperones were previously described to interact with nascent imported proteins, also including CPN60 (Tsugeki and Nishimura, 1993, Madueno et al., 1993, Lubben et al., 1989). Recently it was shown in moss, that the thylakoid transmembrane protein Plastidic type I signal peptidase 1 (Plsp1) utilizes the CPN60 complex for post-translational targeting to thylakoids. It was postulated that the chaperonin captures and releases Plsp1 during targeting (Klasek et al., 2020). We could not identify the Chlamydomonas homolog of Plsp1 (Cre07.g344350) as CPN60 substrate but several imported and integral thylakoid membrane proteins were present in the CPN20/CPN60A substrate sets, including Light Harvesting Complex proteins (LHC) and the photosystem I subunits PsaF and PsaG (Table1). This is consistent with the phenotype of a rice *OsCpn60α1* mutant that displays reduced levels of several photosynthesis related proteins, including PsaG and LHCs (Wu et al., 2020). Thus, CPN60 with its co-chaperonin may also serve in the targeting of thylakoid proteins in Chlamydomonas.

Besides imported proteins, several chloroplast-encoded proteins were amongst the high-confidence substrates, namely RbcL, AtpA, AtpB, ClpP, ChlB, ChlN, PsbB, PsbC, and eight proteins from the 70S translation apparatus (Table 1). We have previously shown, that CPN60 interacts with translating chloroplast ribosomes in a puromycin-sensitive manner, suggesting that the chaperonin facilitates co-translational folding of nascent polypeptides during their synthesis at chloroplast ribosomes (Westrich et al., 2021, Ries et al., 2020). CPN60 binding may start downstream of the ribosome-associated chaperone trigger factor, which may also bind the nascent polypeptides of RbcL and AtpB (Rohr et al., 2019). This agrees with findings in *E. coli*, where the chaperonin cooperates with trigger factor (reviewed in Balchin et al., 2016, Hayer-Hartl et al., 2016) and a small fraction of GroEL binds co-translationally to substrates (Ying et al., 2005). In the cytosol, 5 to 10% of all nascent polypeptides appear to require folding assistance by the CCT/TRiC machinery in a co-translational context (Albanese et al., 2006, Yam et al., 2008). In summary, the chloroplast chaperonin machinery appears to bind nascent polypeptides early upon synthesis and presents a major step for folding prior to complex assembly, at least for Rubisco and the CF_1_ core (Vitlin Gruber and Feiz, 2018, Aigner et al., 2017, Wietrzynski et al., 2021, Mao et al., 2015).

### A special function of the co-chaperonin complex apart from folding

We conducted competition co-IP experiments against the chaperonin complex (i.e.CPN60A) and the lid (i.e. CPN20), since there is accumulating evidence in literature that the co-chaperonin exhibits functions in chloroplasts, which are not directly associated with encapsulating substrates for folding via the CPN60 complex (Zhang et al., 2014, Kuo et al., 2013, Trösch et al., 2015, Zhang et al., 2013). In fact, quantitative immunoblotting indicated that CPN20 accumulates at higher levels when compared to CPN60 (0.17% and 0.1% of the cellular proteome, Figure S2). When considering the molecular mass (20.46 kDa and 58.13 kDa) and the presence of two CPN20 in the lid complex and three CPN60A molecules per heptameric chaperonin ring (Tsai et al., 2012, Zhao et al., 2019), we estimate a seven-fold excess of the lid compared to the chaperonin. In our CPN20 co-IPs, we specifically enriched the almost complete set of catalytically active and inactive subunits of the ClpP protease complex (Figure 3d). Interestingly, CPN20 was previously found to interact with ClpP protease subunits and might even directly or indirectly bind to the ClpC machinery (Kim et al., 2015, Rei Liao et al., 2022). Recently, it has been shown that the CPN23/CPN20/CPN11 lid complex associates with the purified ClpP protease complex, and that the co-chaperonin negatively affects its catalytic activity (Wang et al., 2021). It was hypothesized that this interaction serves to reduce the proteolytic capacity of ClpP and folding via CPN60 at the same time, since the lid is prevented from encapsulating substrates within the CPN60 cavity. We observed an increased interaction between the ClpP machinery and the lid under heat stress conditions. Hence, it might be possible that proteolysis is repressed via the lid complex under these conditions, potentially shifting the equilibrium towards the refolding of denatured proteins if lid complexes are present in excess. In this scenario, the chaperonin might also be shifted to the binding of unfolded proteins to prevent aggregation. Alternatively, the lid may help targeting aberrant substrates for proteolytic cleavage for quality control, however, direct binding of substrates to GroES is rarely observed (Wang et al., 2021, Horwich and Fenton, 2020). The interactions between the co-chaperonin lid and the ClpP protease complex (Wang et al., 2021 and herein) will shed a new light on protein quality pathways of the organelle.

## EXPERIMENTAL PROCEDURES

### Algae strains and growth conditions

The *Chlamydomonas reinhardtii* strain CW15(2)/CF185 (Schroda et al., 1999) was used for all experiments. All cultures were grown mixotrophically in TAP medium (Kropat et al., 2011) on rotary shakers under constant illumination at 80 μmol m^-2^s^-1^ photons (MASTER LEDtube HF 1200 mm UO 16W830/840 T8, Philips) at 25°C. For all experiments, cells were harvested in mid-logarithmic growth phase. For the heat shock experiments cells were resuspended in pre-warmed medium and incubated 30 min at 39°C under the same light regime as before. For individual biological replicates, cells were grown under the same growth conditions, for several days in separate flasks. Endogenous tagging of the plastid encoded ClpP subunit was achieved by homologous recombination via the pUCatpXaadA cassette (Goldschmidt-Clermont, 1991), adding the sequence encoding for a single hemagglutinin (HA) tag 5’-GGTTCATATCCATATGATGTTCCAGATTATGCTTAA-3’, downstream of the *clpP* coding sequence.

### Antibody purification

For the purification of CPN60A and CPN20 antibodies, respective epitopes were covalently coupled to NHS-activated sepharose fast flow beads (Cytiva). Antisera and beads were incubated over night at 4°C, washed with at least 10 column volumes of phosphate-buffered saline (PBS) and brought to pH 8 with 50 mM Tris pH 8, 150 mM NaCl. Antibodies were eluted with pH shock by two rounds of 30 s incubations in 100 mM Glycine pH 2.5, 150 mM NaCl. Upon, elution, pH was directly neutralized by addition of Tris pH 8.8 and buffer was exchanged by dialysis against PBS, containing 10% v/v glycerol.

### Expression of the competitor proteins

For heterologous expression of the competitor proteins in *E. coli*, the DNA sequence encoding for the mature protein of CPN20 (Cre08.g358562), lacking the N-terminal 22 amino acids, was PCR amplified from cDNA clone AV639302 (Kazusa) with oligos 5’-GCCAGGATCCGGAGAATTTATACTTCCAGGGTGCTACCCCCGTGCCCAAG-3’ and 5’-CGCGCGAAGCTTTTACGAGAGCTGGGCCAGG-3’ and cloned with BamHI and HindIII into petDUET (Novagen), giving pFW21. For CPN60A (Cre04.g231222), the sequence coding for the mature protein, lacking the N-terminal 33 amino acids, was PCR amplified from cDNA clone AV640061 (Kazusa), with oligos 5’-GGTGGTTGCTCTTCCAACGCTGACGCTAAGGAGATTGTG-3’ and 5-GGTGGCATATGTTAGATGGTCATGCCGGAGG-3’ and cloned with SapI and NdeI into Tyb21 (NEB), giving pFW38. *E. coli* ER2566 strain (NEB) were grown in M9 minimal medium with ^15^NH_4_Cl as sole nitrogen source as described earlier (Hammel et al., 2018). The CPN60A construct was purified via Chitin-affinity chromatography according to the manufacturer’s instructions (NEB), hexa-histidine-tagged CPN20 was purified with nickel-affinity chromatography under denaturing conditions (Hammel et al., 2018). CPN60A was dialysed into HKM buffer (50 mM Hepes pH 8.0; 25 mM KCl; 25 mM MgCl_2_), CPN20 was dialysed stepwise from 6 M to 4 M, 2 M, 1 M urea in HKM buffer.

### Competition co-immunoprecipitation

For immunoprecipitation, Chlamydomonas cells were harvested by rapidly cooling the culture to below 6°C with -80°C cold permanent ice cubes along with the incubation of 100 μg/mL chloramphenicol (CAP) and centrifugation for 2 min at 4000 g at 4°C. The cell pellet was washed in HKM buffer containing 100 μg/mL CAP and snap frozen as a cell pellet. Cells were lysed in HKM buffer containing 100 μg/mL CAP, 200 μg/μL Heparin, Protease-Inhibitor, 2 mM DSP, 10 mM glucose and 20 U/mL hexokinase (Roche) with the Avestin B-15 at ∼1000 bar homogenizing pressure with 2 cycles. After the lysis 10 mM ADP was added to the lysates and crosslinking was quenched after 30 min of incubation with 100 mM Tris-HCl at pH 8 for 10 min. Protein concentration was determined by Bradford assay (Carl Roth) and Triton X-100 was added to 1%. The lysates were cleared by centrifugation at 20,000 g for 15 min at 4°C and incubated for 1 h with the respective antibody immobilized in Protein-A beads (Amintra) (Willmund and Schroda, 2005). The beads were washed three times with 1% Tween-20 containing buffer (50 mM Hepes pH 8.0; 25 mM KCl; 25 mM MgCl_2_; 100 μg/mL CAP) and three times in the same buffer with 0.1% Tween-20. For HA-affinity purifications from the ClpP-HA strains and CC1690 (recipient strain) at 25 and 39°C the co-IP procedure was modified according to (Westrich et al., 2021). The bound proteins were eluted with sample buffer (125 mM Tris-HCl, pH 6.8, 20% v/v glycerol, 4% w/v SDS and 0.005% bromophenol blue) lacking DTT by boiling for 1 min. To revert the cross-link, 50 mM DTT was added to the eluates and boiled again for 5 min.

For competition co-IP, detailed protocols are included in our deposited arc (see Data availability), including information on the pre-experiment. Cell lysates from three independent cultures were split into 6 equal fractions and competitor protein was added at concentrations that were 0.25, 0.5, 1, 2.5, 5 and 10-fold relative to the target protein before adding the antibody-coupled beads. For mass spectrometric measurements, samples were loaded on a 10% SDS-PAGE and run 1 cm into the separating gel. Protein bands were stained with colloidal Coomassie G and gel pieces of 10 mm^2^ were used for in-gel digestion. Protein digest, and mass spectrometry was performed as described earlier (Westrich et al., 2021).

Extraction of ion chromatograms and the identification and quantification of labeled (^15^N) and unlabeled peptides was performed using ProteomIQon software suite v0.0.2 (https://doi.org/10.5281/zenodo.6335068). Default parameters were used with methionine oxidation and N-term acetylation as additional variable modifications and data were searched against the reference Chlamydomonas genome (JGI5.5), including plastid encoded sequences.

Peptide quantification was obtained by aggregating of different charge states for each peptide and averaging over the technical replicates. Peptides identified in less than 5 of 6 samples of the target protein concentration series were removed from the dataset. After averaging experiments for individual biological replicates, protein inference was performed using the average over all respective peptides. Statistical analysis was performed using the FSharp.Stats library v0.4.4 (https://doi.org/10.5281/zenodo.6337056). Correlation of target protein was calculated against all proteins in the concentration series using weights based on inverse standard deviation of each datapoint over biological replicates. Correlation *p*-values were obtained using jackknife cross validation and corrected for multiple testing by estimating *q*-values (Storey and Tibshirani, 2003). For each experiment, proteins with correlation values below 0.7 and a *q*-value above 0.05 were filtered out.

Protein properties except aggregation propensities were computed using the BioFSharp library v2.0.0 (https://doi.org/10.5281/zenodo.6335372). Aggregation propensity was computed using Tango tool v2.3 (Rousseau et al., 2006). Here, the pH was set to 7.8, ionic strength was set to 0.26 and organic solvent percentage was set to 0. The temperature was set dependent on the specific experiment, 298K or 312K.

### Miscellaneous

SDS–PAGE, semi-dry blotting and immunodetection were carried out as described before (Willmund and Schroda, 2005). Immunoblots were captured with the Intas ChemoStar 6.0. The abundance of the target proteins was determined by quantitative immunoblotting using the recombinant epitope as a quantification standard (Rohr et al., 2019).

## ACKNOWLEDGEMENTS

We thank Karin Gries for technical assistance with protein and antibody purification and Cuimin Liu for critical discussion of the data. This work was supported by the Carl-Zeiss fellowship to F.R., the Deutsche Forschungsgemeinschaft grant TRR175 A05, C02, D02 to F.W. M.S., T.M. respectively, and the Forschungsschwerpunkt BioComp to T.M., M.S., F.W.

## AUTHOR CONTRIBUTIONS

F.R. designed and performed the experiments and wrote parts of the text, C.H. generated the ClpP-HA strain. H.L.W. and T.M. evaluated mass spectrometry data. F.S. and M.S. provided mass spectrometry data. F.W. designed the project, evaluated data, and wrote the manuscript.

## DATA AVAILABILITY

All gene accession codes are given in the Supporting Dataset and are in accordance with the latest Chlamydomonas genome v6.1 annotation (https://phytozome-next.jgi.doe.gov) or the GenBank/EMBL data libraries. The mass spectrometry data are accessible via the Dataplant consortium (https://git.nfdi4plants.org/weil/CPN2060_CoIP).

## SUPPORTING INFORMATION

Additional Supporting Information is found in the online version of this article.

### Supporting Dataset

Spreadsheet of proteomic data from all replicates, subcellular localization, functional annotation, correlation *r*-values and *q*-values.

## SUPPORTING INFORMATION

**Figure S1:**
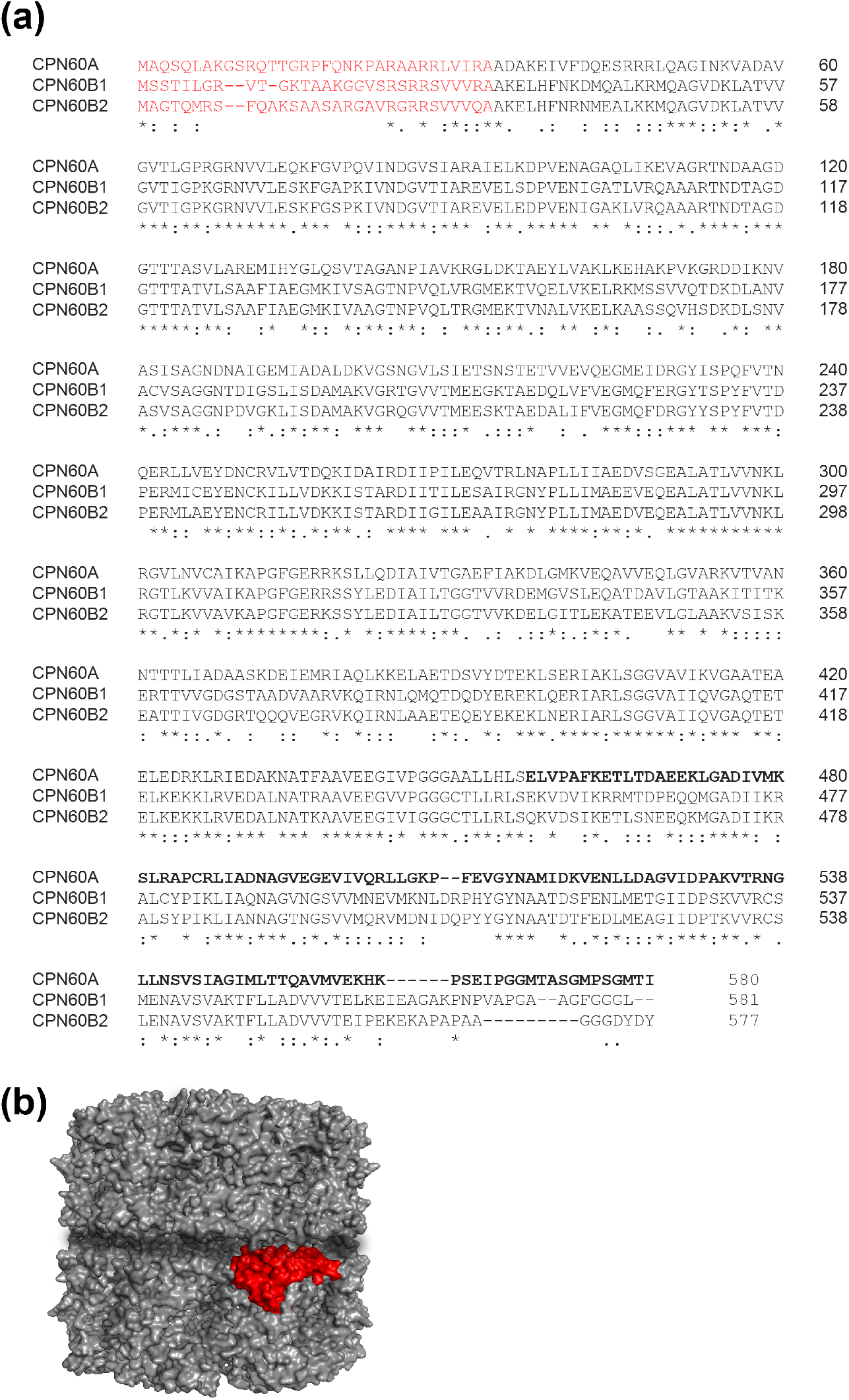
Sequence alignment and model of the chloroplast CPN60 subunits in Chlamydomonas. (**a**) Alignment of the chloroplast CPN60 amino acid sequences from *Chlamydomonas reinhardtii*: CPN60A (Cre04.g231222), CPN60B1 (Cre17.g741450) and CPN60B2 (Cre07.g339150). Predicted chloroplast transit peptides are marked in red. The C-terminal 124 amino acids, which were used as antigen for CPN60A antibody generation, are highlighted in bold letters. Sequences were aligned by ClustalOmega. (**b**) Structural model of the Chlamydomonas CPN60 complex with the CPN60A epitope marked in red, based on PDB 5cdi (Zhang et al., 2016).

**Figure S2:**
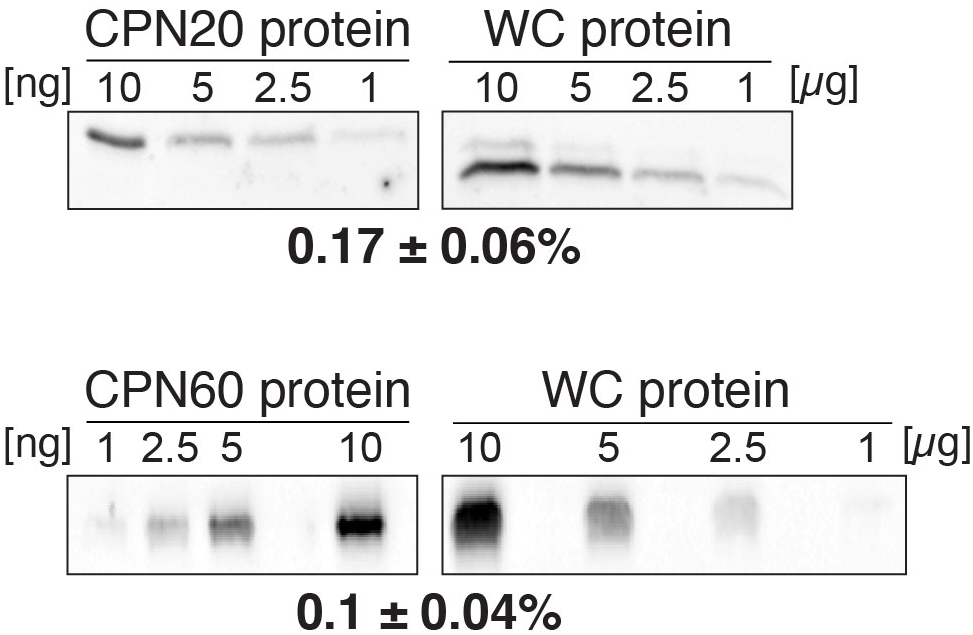
Quantification of protein concentrations in cell lysates. Quantitative immunoblots of CPN20 (top) and CPN60A (bottom) of lysates of Chlamydomonas cells. Lanes on the left contain a defined concentration range of heterologously produced, purified proteins. Lanes on the right contain defined amounts of whole-cell Chlamydomonas protein. Percentages of CPN20 and CPN60A proteins were calculated on three biological replicates.

**Figure S3:**
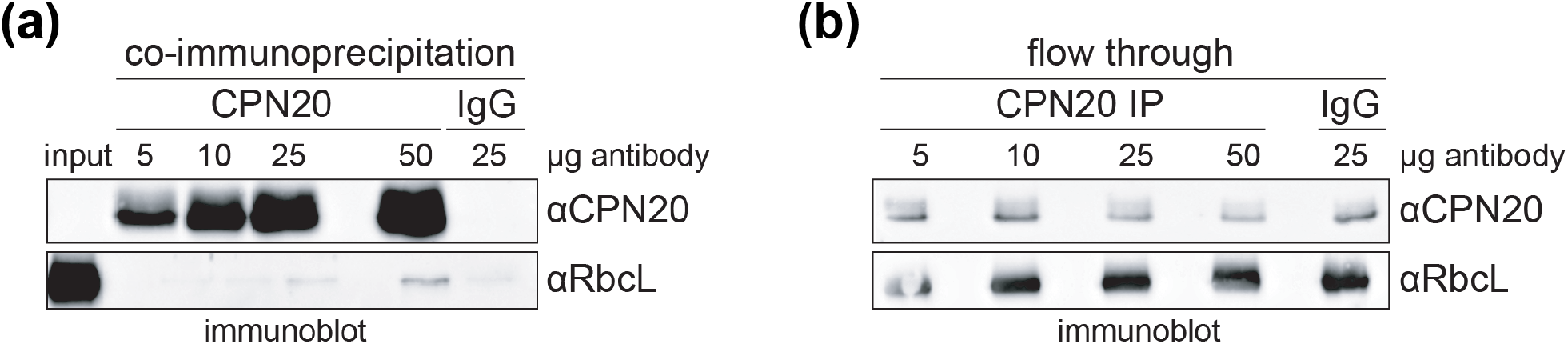
Test of target protein depletion by co-immunoprecipitation. (**a**) CPN20 IPs with increasing amounts of purified α-CPN20 antibody and with unspecific rabbit IgGs from 5×10^8^ Chlamydomonas cells per sample. Immunoblot targeted against the IP eluates against CPN20 and the known chaperonin substrate RbcL. (**b**) Immunoblot of the flow through samples (lysate after immune-depletion), assayed for the remaining amounts of CPN20 and RbcL. IgG corresponds to unspecific and purified antibodies.

**Figure S4:**
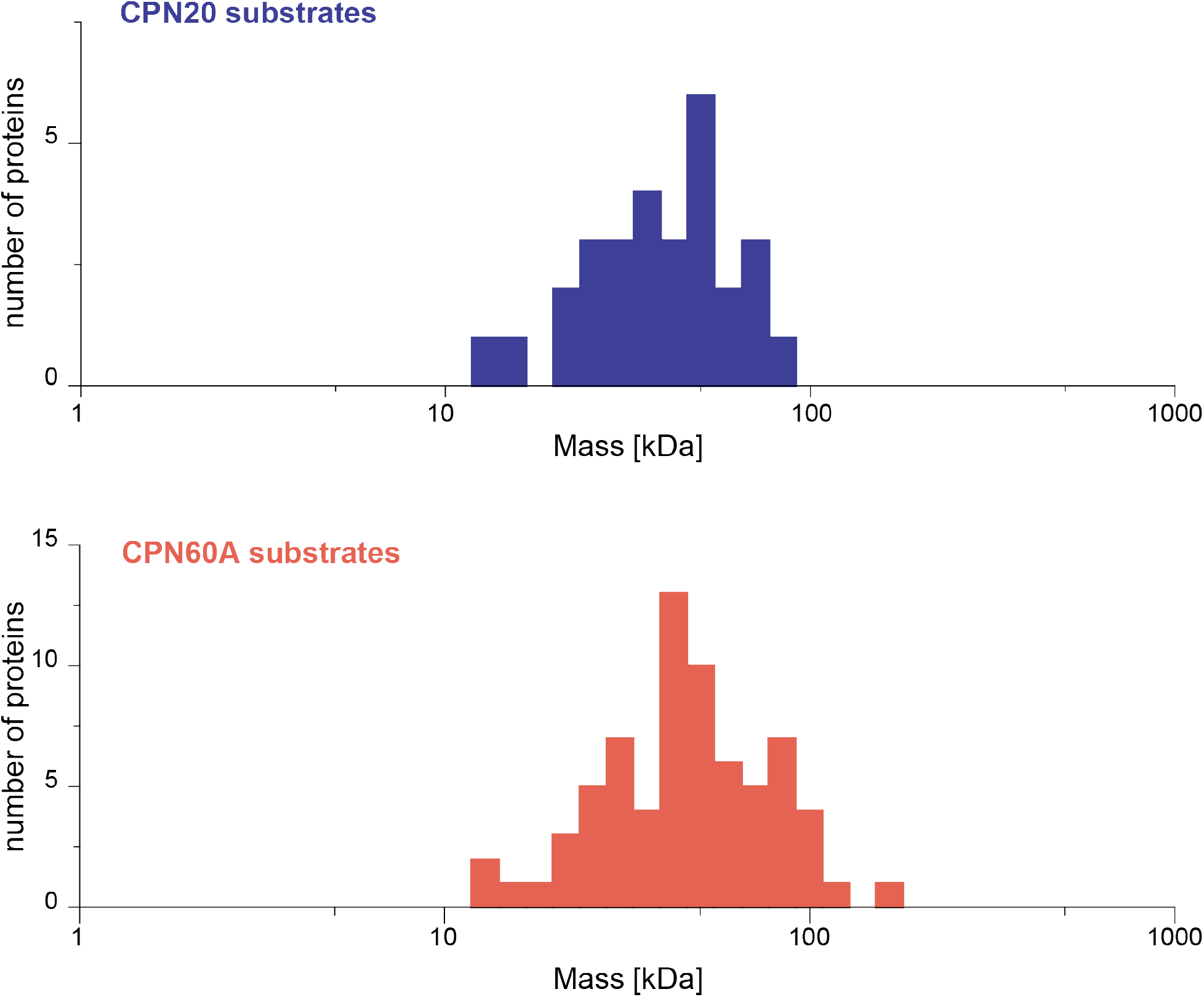
Size distribution of chaperonin substrates. Histogram of molecular mass (kDa) for the putative chaperonin substrates in the CPN20 (blue) and CPN60A (red) datasets. Molecular mass was plotted for the proteins with correlation *r*-values ≥ 0.7 and *q*-values < 0.05, excluding known interactors such as the chaperonin complex partner proteins and the ClpP complex (CPN20 only).

**Figure S5:**
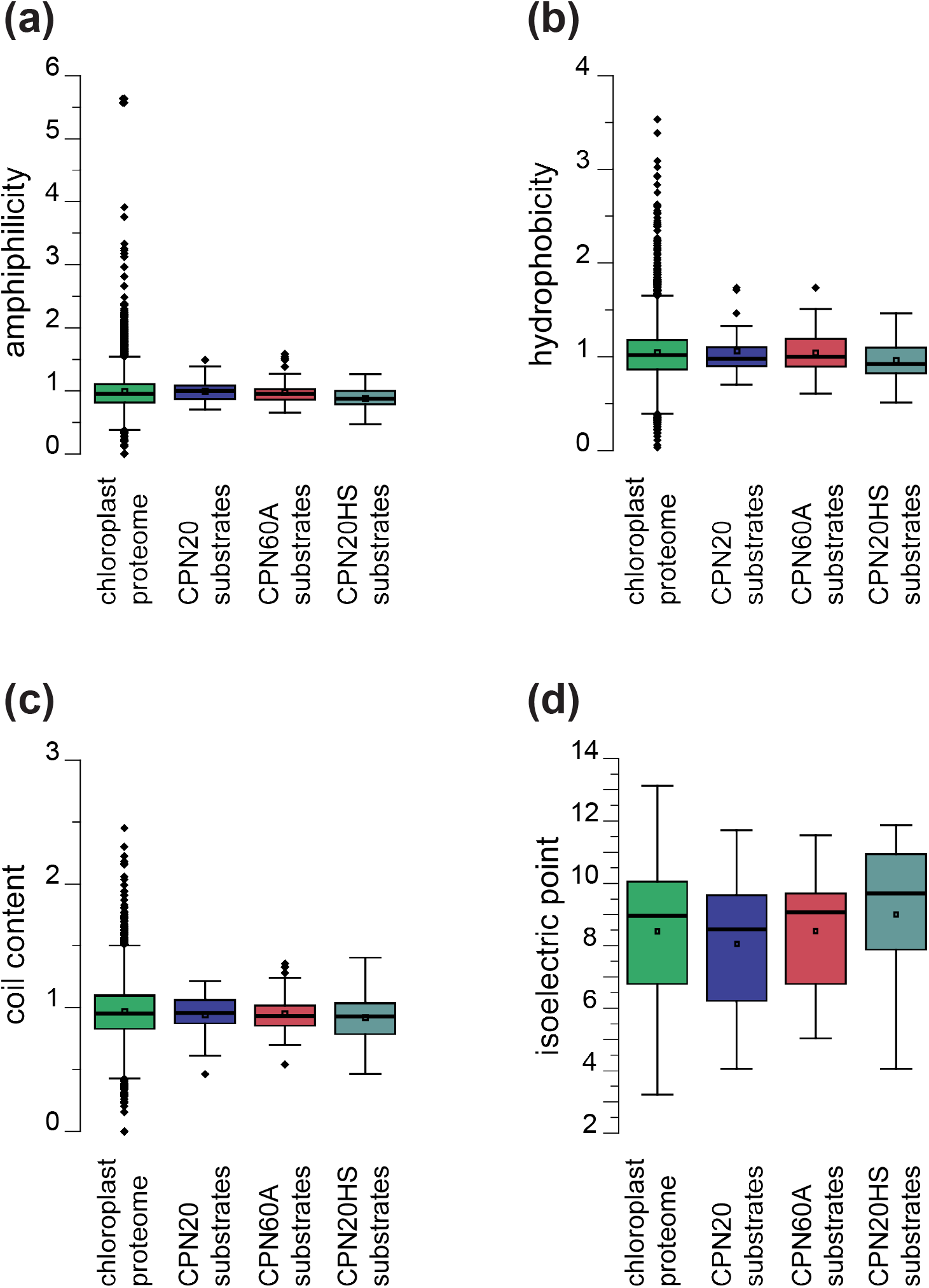
Physicochemical protein properties of chaperonin substrates. Additional properties analyzed, accompanying Figure 5, comparing all proteins with a chloroplast localization (green), CPN20 (blue) and CPN60A (red) substrates and of CPN20 after a shift to 39°C for 30 min (turquoise). (**a**) Boxplot showing distribution of amphiphilic values for proteins in the dataset. (**b**) Boxplot comparing hydrophobicity of the amino acid sequence. (**c**) Boxplot comparing content of coiled secondary structures. (**d**) Boxplot comparing distribution of isoelectric points for the analyzed proteins. All values were tested for significance by students *t*-test assuming equal variance. No *p*-values are given if non-significant.

